# selscape: A Snakemake Workflow for Investigating Genomic Landscapes of Natural Selection

**DOI:** 10.64898/2026.02.28.708717

**Authors:** Simon Chen, Xin Huang

## Abstract

Analyzing natural selection is a central task in evolutionary genomics, yet applying multiple tools across populations in a reproducible and scalable manner is often complicated by heterogeneous input formats, parameter settings, and tool dependencies. Here, we present selscape, a Snakemake workflow that automates end-to-end genome-wide selection analysis—from input preparation and statistic calculation to functional annotation, downstream visualization, and summary reporting. We demonstrate selscape on high-coverage genomes from the 1000 Genomes Project, illustrating how the workflow enables efficient, large-scale analyses and streamlined comparisons across populations. By unifying diverse tools with Snakemake, selscape lowers the barrier to robust genome-wide analyses and provides a flexible framework for future extensions and integration with complementary population genetic analyses.

---

Natural selection shapes genetic diversity in various species by acting on heritable variants with differential fitness (Nei 1987; Crow and Kimura 2009; Felsenstein 2019). With expanding genomic resources and an increasingly rich methodological toolkit, researchers can detect and quantify signatures of natural selection within and between populations, linking evolutionary history to functional genomic variation (Sackton 2020; Bitarello et al. 2023; Panigrahi et al. 2023). However, existing methods are often scattered across disparate software packages with inconsistent input and output formats, making it difficult to run multiple selection analyses in a standardized, reproducible manner. Meanwhile, the growing adoption of workflow management system, such as Snakemake (Köster and Rahmann 2012; Mölder et al. 2021), has provided a practical path to package and distribute complete analysis recipes rather than isolated tools.

Snakemake defines analyses as a set of rules connected by input–output dependencies. It manages software dependencies via Conda, enabling convenient installation and consistent environments for heterogeneous tools, and scales from local machines to high-performance computing clusters while supporting reproducible analyses. This makes it well suited for orchestrating multiple tools into a single workflow that is portable across diverse computing platforms. Recently, Snakemake workflows have been developed for different tasks in population genomics, including variant calling, basic population genomic analyses, and simulation-based inference (Mirchandani et al. 2024; Min et al. 2025; Nolen 2025); but to our knowledge, no comparable workflow currently exists for whole-genome selection analyses. Here, we present selscape (version 1.0.0), a Snakemake workflow that automates a suite of complementary selection analyses with standardized preprocessing, parameterized execution, and unified outputs and summary reports.

In selscape (Figure 1), scikit-allel, selscan, BetaScan, and dadi-cli are integrated to run complementary analyses targeting different modes of natural selection. Tajima’s *D* is computed with scikit-allel to capture allele-frequency-spectrum deviations consistent with positive or balancing selection (Tajima 1989; Miles et al. 2024). Haplotype-based statistics for recent positive selection, including iHS, nSL, XP-EHH, and XP-nSL, are estimated with selscan (Voight et al. 2006; Sabeti et al. 2007; Ferrer-Admetlla et al. 2014; Szpiech and Hernandez 2014; Szpiech et al. 2021; Szpiech 2024; Rahman et al. 2025a, 2025b). Long-term balancing selection is also assessed using BetaScan (Siewert and Voight 2017; Siewert and Voight 2020). The distribution of fitness effects (DFE) is inferred using dadi through its command-line interface—dadi-cli (Gutenkunst et al. 2009; Huang et al. 2023). Upstream of these selection analyses, selscape performs standardized preprocessing with BCFtools and PLINK for data processing and variant filtering (Chang et al. 2015; Danecek et al. 2021). Downstream, selscape supports biological interpretation through Gene Ontology (GO) enrichment tests of outlier variants using Gowinda, which accounts for biases due to gene length (Kofler and Schlötterer 2012).

**Figure 1.**
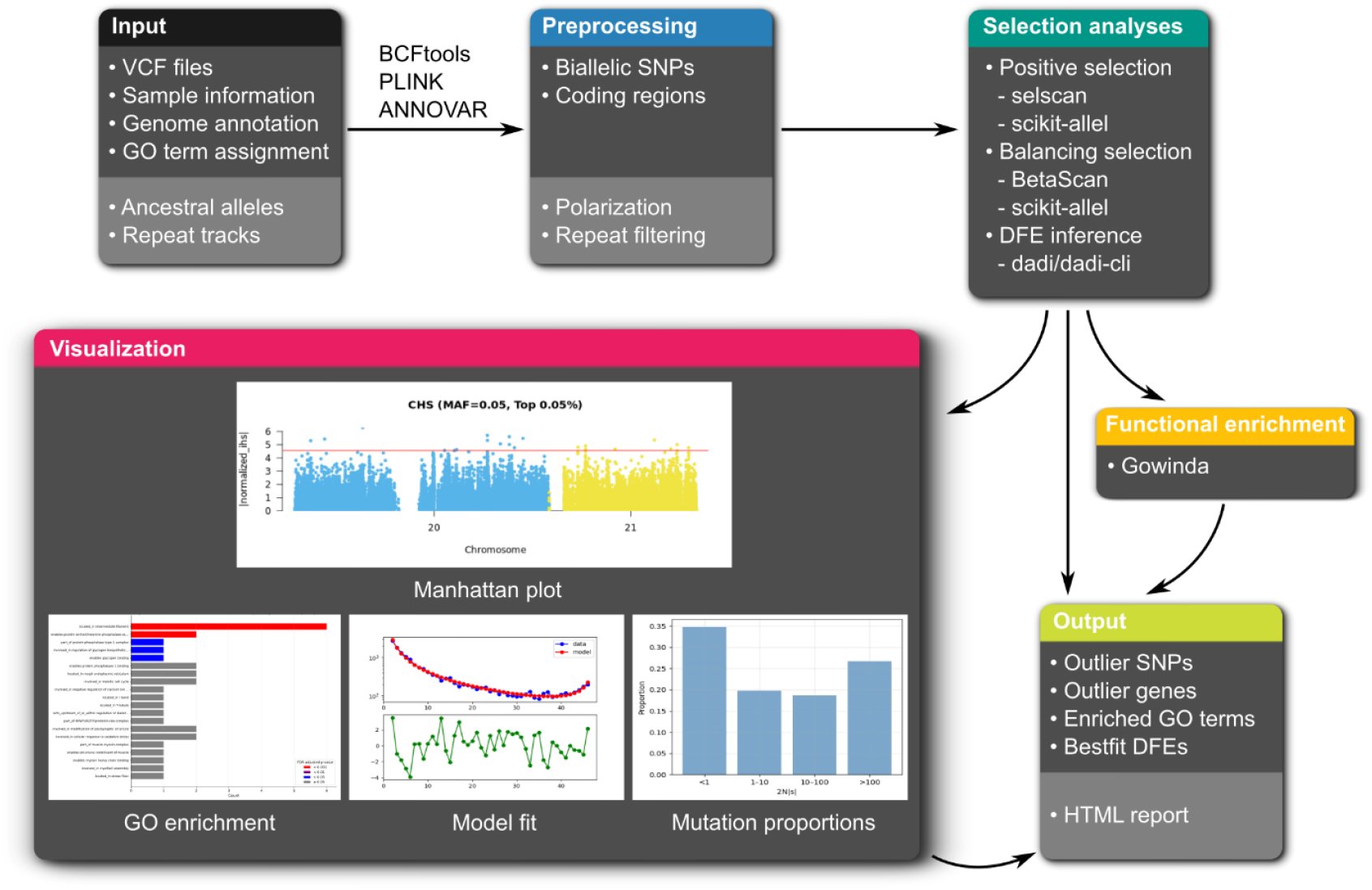
Overview of selscape. The workflow consists of multiple modular components. Required inputs include VCF files containing genotypes, sample information defining population assignments, a genome annotation file, and a gene ontology (GO) assignment file. Optional inputs include ancestral allele information for polarization and repeat tracks for filtering repetitive regions. Core preprocessing steps include extracting biallelic single nucleotide polymorphisms (SNPs) and stratifying coding-region sites into synonymous and nonsynonymous classes. The processed data are then used for 3 types of selection analyses: detection of positive selection using selscan and scikit-allel, detection of balancing selection using BetaScan and scikit-allel, and inference of the distribution of fitness effects (DFE) using dadi via dadi-cli. Outliers from positive and balancing selection analyses are passed to Gowinda for GO enrichment testing. The workflow generates multiple plots, including Manhattan plots for selection statistics, model fit plots for demographic and DFE inference, plots for the proportions of deleterious mutations with different selection coefficients, and plots for GO enrichment. The workflow also produces tables summarizing outlier SNPs and outlier genes, GO enrichment results, and best fitting DFE parameters with their estimated 95% confidence intervals. These tables and plots can optionally be compiled into an interactive HTML report.

In addition, ANNOVAR is used to stratify variants into synonymous and nonsynonymous classes for DFE inference and to annotate outlier variants and map them to genes for downstream interpretation (Wang et al. 2010). Results are visualized using qqman for Manhattan plots (Turner 2018), dadi for allele frequency spectra and residuals between the inferred and observed spectra, and matplotlib for summary plots of deleterious-mutation proportions and enriched GO terms. Finally, an interactive HTML report containing tables and plots can be generated using Snakemake’s built-in reporting functionality. Components of selscape have been used in multiple studies to analyze ancient and modern human and non-human great ape genomes for natural selection (Gelabert et al. 2025; Hackl and Huang 2025; Huang et al. 2025a).

They were also applied in the selective sweep challenges of the Genomic History Inference Strategies Tournament 2025 (Struck et al. 2025; Chen and Huang 2025).

To demonstrate selscape in practice, we used it to analyze 2,504 high-coverage genomes from 26 world-wide human populations in the 1000 Genomes Project (Byrska-Bishop et al. 2022). We scanned these genomes for signatures of positive selection with single-population (iHS, nSL, Tajima’s *D*) and cross-population statistics (XP-EHH, XP-nSL, ΔTajima’s *D*), and for balancing selection with summary statistics (*β*^(1)^ and Tajima’s *D*). As expected, our scans recapitulate well-established signals of positive selection at genes associated with human pigmentation, such as *SLC24A5, SLC45A2*, and *OCA2* (He et al. 2015; Huang et al. 2021; He et al. 2025). In addition, we recover classic signatures of balancing selection in the Human Leukocyte Antigen (HLA) region. Figure 2 displays the genome-wide results of positive selection in the CHS population based on single-population statistics.

**Figure 2.**
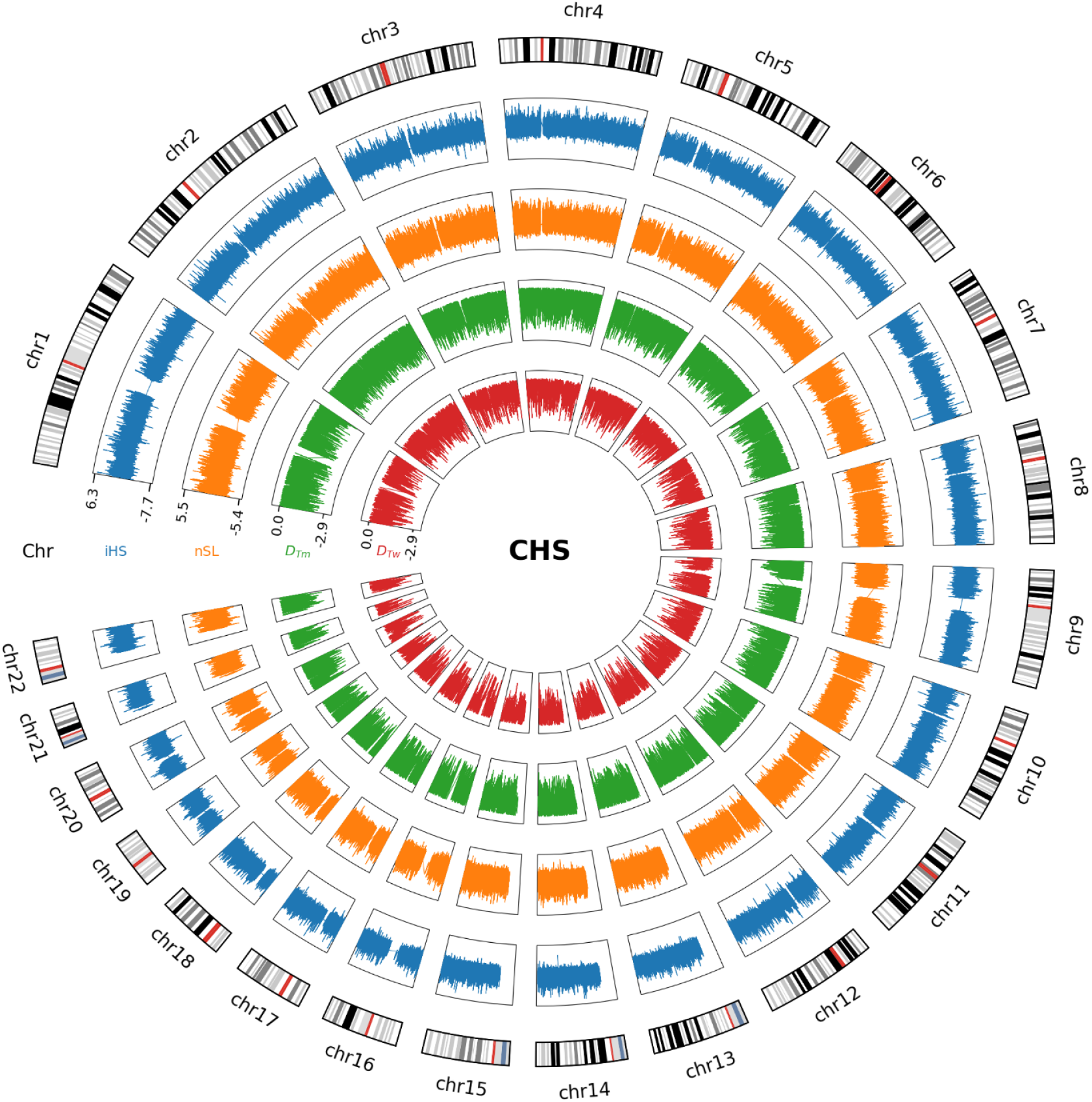
Circos plot for genome-wide positive selection scan in the CHS population with single-population statistics. From the outer to the inner track: iHS, integrated haplotype score; nSL, number of segregating sites by length; *D*_*Tm*_, Tajima’s *D* computed in sliding windows with a fixed number of SNPs, and *D*_*Tw*_, Tajima’s *D* computed in sliding windows with a fixed number of base pairs.

We also inferred lognormal DFEs for these populations and estimated confidence intervals (CIs) using a Godambe approach (Coffman et al. 2016). Relative to estimates based on 1000 Genomes Project Phase 3 data (The 1000 Genomes Project Consortium 2015; Huang et al. 2023), our results yield comparable CIs for the mean parameter (*μ*), but substantially narrower CIs for the standard deviation (*σ*) (Figure 3). In comparison with DFEs inferred from a curated great ape genomes dataset (Han et al. 2025; Huang et al. 2025a), our CIs broadly overlap those of non-human great apes, suggesting that these DFE parameters may be largely conserved across great apes. This pattern is consistent with a previous study that assumed gamma DFEs (Castellano et al. 2019).

**Figure 3.**
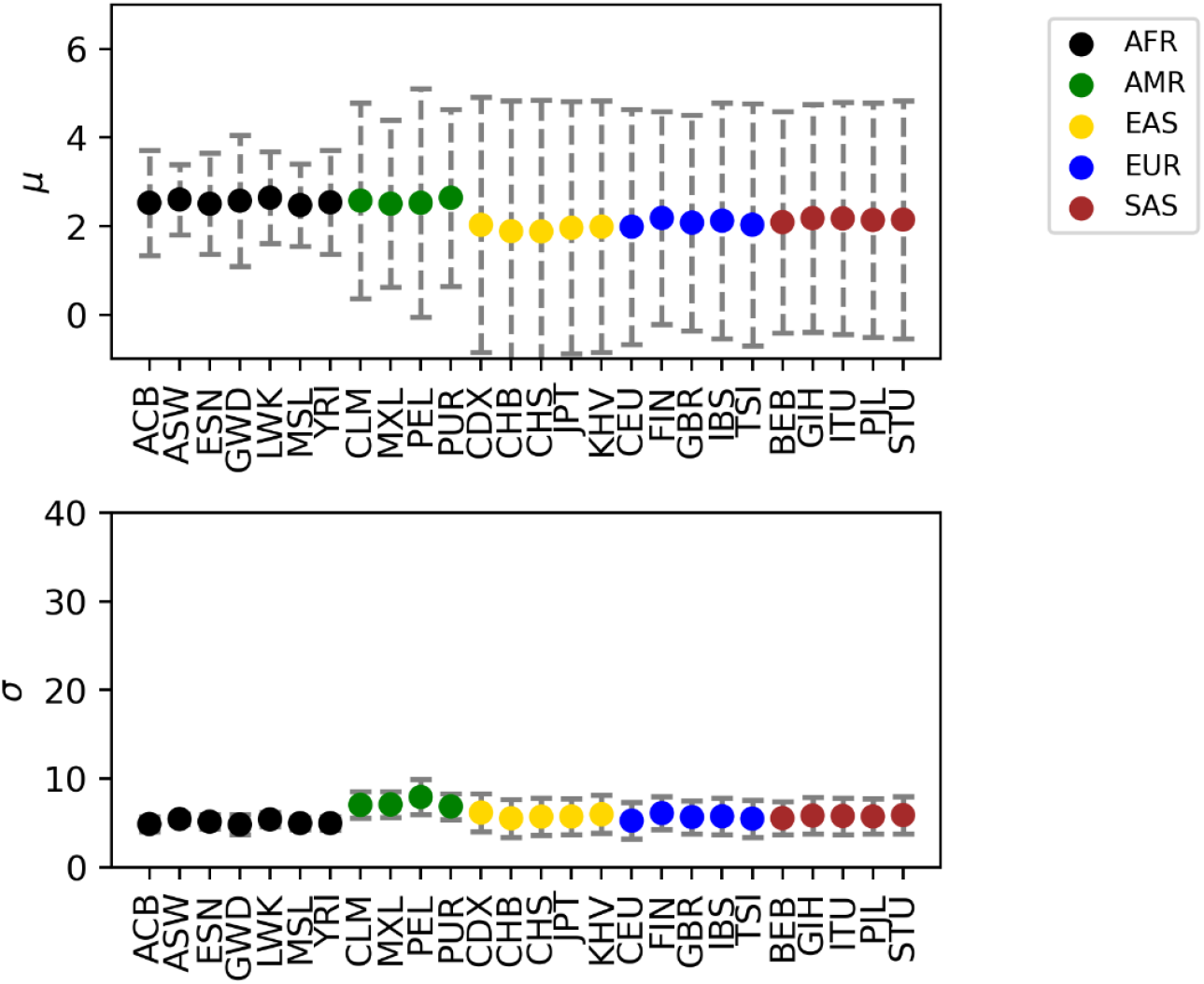
95% Confidence intervals for the estimated DFE parameters assuming a lognormal distribution. *μ* and *σ* are the mean and standard deviation of the log of the population-scaled selection coefficient, respectively. AFR, African populations; AMR, American populations; EAS, East Asia populations; EUR, European populations; SAS, South Asia populations; ACB, African Caribbean in Barbados; ASW, African Ancestry in Southwest US; ESN, Esan in Nigeria; GWD, Gambian in Western Division, The Gambia; LWK, Luhya in Webuye, Kenya; MSL, Mende in Sierra Leone; YRI, Yoruba in Ibadan, Nigeria; CLM, Colombian in Medellin, Colombia; MXL, Mexican Ancestry in Los Angeles, California; PEL, Peruvian in Lima, Peru; PUR, Puerto Rican in Puerto Rico; CDX, Chinese Dai in Xishuangbanna, China; CHB, Han Chinese in Beijing, China; CHS, Han Chinese South; JPT, Japanese in Tokyo, Japan; KHV, Kinh in Ho Chi Minh City, Vietnam; CEU, Utah residents (CEPH) with Northern and Western European ancestry; FIN, Finnish in Finland; GBR, British in England and Scotland; IBS, Iberian populations in Spain; TSI, Toscani in Italia; BEB, Bengali in Bangladesh; GIH, Gujarati Indian in Houston, TX; ITU, Indian Telugu in the UK; PJL, Punjabi in Lahore, Pakistan; STU, Sri Lankan Tamil in the UK.

In conclusion, selscape provides an automated and scalable workflow for investigating genomic landscapes of natural selection. As machine learning-based methods in population genetics continue to expand (Schrider and Kern 2018; Huang et al. 2024), selscape can be readily extended to incorporate and interface with these approaches. Other population genetic tasks that involve multiple tools, such as the detection of introgressed fragments (Huang et al. 2025b), can likewise be integrated and managed within a unified Snakemake framework.

## Acknowledgements

The authors thank the Life Science Compute Cluster at the University of Vienna and the Multi-Site Computer Austria of Austrian Scientific Computing for providing computing resources.

## Author contributions

X.H. designed the study. S.C. and X.H. implemented selscape, analyzed the data and wrote the manuscript.

## Data availability

The high-coverage data from the 1000 Genomes Project can be found in https://ftp.1000genomes.ebi.ac.uk/vol1/ftp/data_collections/1000G_2504_high_coverage/working/20190425_NYGC_GATK/. The ancestral allele information based on the hg38 reference genome can be found in https://ftp.ensembl.org/pub/release-115/fasta/ancestral_alleles/homo_sapiens_ancestor_GRCh38.tar.gz. The RepeatMasker Track from the UCSC Genome Browser can be found in https://hgdownload.soe.ucsc.edu/goldenPath/hg38/database/rmsk.txt.gz. The Segmental Dups Track from the UCSC Genome Browser can be found in https://hgdownload.soe.ucsc.edu/goldenPath/hg38/database/genomicSuperDups.txt.gz. The Simple Repeats Track from the UCSC Genome Browser can be found in https://hgdownload.soe.ucsc.edu/goldenPath/hg38/database/simpleRepeat.txt.gz. Genome annotation can be found in https://ftp.ncbi.nih.gov/genomes/refseq/vertebrate_mammalian/Homo_sapiens/annotation_releases/GCF_000001405.40-RS_2024_08/GCF_000001405.40_GRCh38.p14_genomic.gtf.gz. GO term assignment can be found in https://ftp.ncbi.nih.gov/gene/DATA/gene2go.gz. BCFtools can be found in https://github.com/samtools/bcftools. PLINK can be found in https://www.cog-genomics.org/plink/. selscan can be found in https://github.com/szpiech/selscan. BetaScan can be found in https://github.com/ksiewert/BetaScan. dadi-cli can be found in https://github.com/xin-huang/dadi-cli. scikit-allel can be found in https://github.com/cggh/scikit-allel. ANNOVAR can be found in https://annovar.openbioinformatics.org/en/latest/. Gowinda can be found in https://sourceforge.net/projects/gowinda/. selscape can be found in Snakemake Workflow Catalog: https://snakemake.github.io/snakemake-workflow-catalog/docs/workflows/xin-huang/selscape.html. The source code of selscape can be found in https://github.com/xin-huang/selscape. The Snakemake workflow for reproducing the analysis can be found in https://github.com/xin-huang/selscape-analysis. All URLs were last accessed on February 28, 2026.

